# Phylogeny and distribution of Y-chromosomal haplotypes in domestic, ancient and wild goats

**DOI:** 10.1101/2020.02.17.952051

**Authors:** Isaäc J. Nijman, Benjamin D. Rosen, Zhuqing Zheng, Yu Jiang, Tristan Cumer, Kevin G. Daly, Valentin A. Bâlteanu, Beate Berger, Thor Blichfeldt, Geert Boink, Sean Carolan, Vlatka Cubric-Curik, Juha Kantanen, Amparo Martínez, Raffaele Mazza, Negar Khayatzadeh, Namshin Kim, Nadjet-Amina Ouchene-Khelifi, Filipe Pereira, Anne da Silva, Mojca Simčič, Johann Sölkner, Alison Sutherland, Johannes Tigchelaar, Econogene Consortium, Paolo Ajmone-Marsan, Daniel G. Bradley, Licia Colli, François Pompanon, Johannes A. Lenstra

**Author notes:** See the list at the end of this paper (Econogene Consortium). **Correspondence:** J.A. Lenstra.

## Abstract

The male-specific part of the Y-chromosome is in mammalian and many other species the longest haplotype that is inherited without recombination. By its paternal transmission it has a small effective population size in species with dominant males. In several species, Y-chromosomal haplotypes are sensitive markers of population history and introgression. Previous studies have identified in domestic goats four major Y-chromosomal haplotypes Y1A, Y1B, Y2A and Y2B with a marked geographic differentiation and several regional variants. In this study we used published whole-genome sequences of 70 male goats from 16 modern breeds, 11 ancient-DNA samples and 29 samples from seven wild goat species. We identified single-copy male-specific SNPs in four scaffolds, containing *SRY, ZFY, DBY* with *SSX3Y* and *UTY,* and *USP9Y* with *UMN2001,* respectively. Phylogenetic analyses indicated haplogroups corresponding to the haplotypes Y1B, Y2A and Y2B, respectively, but Y1A was split into Y1AA and Y1AB. All haplogroups were detected in ancient DNA samples from southeast Europe and, with the exception of Y1AB, in the bezoar goat, which is the wild ancestor of the domestic goats. Combining these data with those of previous studies and with genotypes obtained by Sanger sequencing or the KASP assay yielded haplogroup distributions for 132 domestic breeds or populations. The phylogeographic differentiation indicated paternal population bottlenecks on all three continents. This possibly occurred during the Neolithic introductions of domestic goats to those continents with a particularly strong influence in Europe along the Danubian route. This study illustrates the power of the Y-chromosomal haplotype for the reconstructing the history of mammalian species with a wide geographic range.

## 1. Introduction

Because of its absence or recombination, the male part of the mammalian Y-chromosomes is by far the longest haplotype that is stably transmitted across generations (Hughes et al., 2015). Together with its inheritance from father to son, it is a highly informative marker for the paternal origin of species, populations or individuals with a much stronger phylogeographic differentiation than observed for mitochondrial or autosomal DNA. The highly informative Y-chromosomal markers are now widely exploited in population-genetic studies of humans (Jobling and Tyler-Smith, 2017; Kivisild, 2017), cattle (Edwards et al., 2011; Xia et al., 2019), horse (Wallner et al., 2017; Wutke et al., 2018; Felkel et al., 2019a) water buffalo (Zhang et al., 2016), sheep (Meadows and Kijas, 2009; Zhang et al., 2014), camel (Felkel et al., 2019b), pigs (Guirao-Rico et al., 2018) and dogs (Natanaelsson et al., 2006; Oetjens et al., 2018).

A preliminary analysis of the Y-chromosomal diversity in European and Turkish goats defined the three haplotypes Y1A, Y1B and Y2, showing strong geographic differentiation (Lenstra and Econogene Consortium, 2005). The same haplotypes were found in goats from Portugal and North-Africa (Pereira et al., 2009), Turkey (Cinar Kul et al., 2015), east and south Asia (Waki et al., 2015), Switzerland and Spain (Vidal et al., 2017) with additional haplotypes Y2B in east Asia, Y2C in Turkish Kilis goats, and Y1B2 as well Y1C mainly in Switzerland (Table 1). However, these haplotypes are based on a low number of SNPs in or near *DBY-1, SRY* and ZFY. Thus, it is not clear if the haplotypes represent major haplogroups or local variants, if other major haplogroups exist, and which of the Y-chromosomal variants already existed in the earlier domestic goats and in their wild ancestor, the bezoar (Amills et al., 2017).

**Table 1.** Haplotypes identified previously or in this study by ABI sequencing (Lenstra, 2005; indicated by ddSeq), whole-genome sequencing, KASP genotyping, SRA BLAST searches. Mutations are relative to haplogroup Y1AB, which in previous studies together with Y1AA was represented by haplotype Y1A. Haplotype Y1C belongs to the Y1B haplogroup. Most of the diagnostic SNPs used here for KASP or SRA searches are within the segments covered by dideoxy sequencing.

| Haplotypes | Diagnostic SNP | Detection | Specificity | Range, breed | Reference |
| --- | --- | --- | --- | --- | --- |
| **Main types** |  |  |  |  |  |
| Y1AA | NW_017189563.1 g.G280574>T | ddSeq, SRA search | Y1AA | Asia<sup>1</sup> | This study |
| Y1AB |  |  |  | Europe, Asia<sup>1</sup> | Lenstra, 2005 |
| Y1B | NW_017189563.1 g.T280306>A | ddSeq, SRA search, KASP | Y1B, Y1C | Europe, North Africa | Lenstra, 2005 |
|  | NW_017189885.1 g.T104701>C | SRA search | Y1B, Y1C | Europe, North Africa | This study |
| Y2A | NW_017189563.1 g.G277537>T | ddSeq, SRA search | Y2A, Y2C, Y2C |  | Lenstra, 2005 |
|  | NW_017189563.1 g.A278523>G | ddSeq, SRA search | Y2A, Y2C, Y2C |  | Lenstra, 2005 |
|  | NW_017189563.1 g.T280046>A | ddSeq | Y2A, Y2C, Y2C |  | Lenstra, 2005 |
|  | NW_017189563.1 g.G280433>A | ddSeq | Y2A, Y2C, Y2C |  | Lenstra, 2005 |
|  | NW_017189685.1 g.A43279>C | ddSeq | Y2A, Y2C, Y2C |  | Lenstra, 2005 |
|  | NW_017189685.1 g.C43464>T | ddSeq | Y2A, Y2C, Y2C |  | Lenstra, 2005 |
|  | NW_017189885.1 g.A11686>G | ddSeq, KASP | Y2A, Y2C, Y2C |  | Lenstra, 2005 |
| Y2B | NW_017189563.1 g.A280113>G | ddSeq, SRA search | Y2B | East, southeast Asia | Waki et al., 2015 |
| **Local types** |  |  |  |  |  |
| Y1AB2 | NW_017189563.1 g.A277871>G | ddSeq | Y1AB2 | Albanian DUK, LIQ; Romanian CAR; German TWZ | This study |
| Y1B2 | NW_017189885.1 g.G12927>A |  | Y1B2 | Switzerland, Italy | Vidal et al., 2017 |
| Y1C | NW_017189563.1 g.A279212>C | ddSeq | Y1C | Swiss PCG, VZC; Italian GAR | Vidal et al., 2017 |
| Y2C | NW_017189885.1 g.A11760>C |  | Y2C |  | Cinar-Kul et al., 2015 |

<sup>1</sup> Y1A is present in Africa and may represent either Y1AA or Y1AB or both

In this study, we use whole-genome sequences (WGSs) to systematically characterize the SNP-level variation in a large part of the single-copy male-specific part of the caprine Y-chromosome. In addition, we determined the Y-chromosomal haplogroups in goats originating from several European, Asian or African countries, in ancient goat DNA samples and in the wild bezoar. We observed a strong phylogeographic structure as the result of paternal bottlenecks during the Neolithic migrations and found evidence for recent introgressions in Asian landraces, which is essential information for breed management and conservation.

## 2. Methods

### 2.1 WGS data, filtering and tree construction

We selected four Y-chromosomal scaffolds carrying single-copy genes without the UMN2303 Y-chromosomal repeat (Table 2). From the Short Read Archive (SRA) a VCF file containing the sequences of these scaffolds in 70 male goats (Table 3), 30 of which were from central and east Asia, and 107 female goats from the same source laboratories. After excluding dense clusters of SNPs, we obtained 5356 SNPs, from which 2350 were hemizygous, were not scored in females, had scores in >95% of the males and had a minor allele frequency of >2% in male domestic goats. Allele-sharing distances were calculated using Plink, and visualized in a Neighbor-Joining tree by using the programs Splitstree (Huson and Bryant, 2006) and Mega (Tamura et al., 2011).

**Table 2.** X- and Y-chromosomal scaffolds for the goat reference genome.

| scaffold | length (bp) | genes |
| --- | --- | --- |
| NW_017189563.1 | 528190 | *SRY* |
| NW_017189618.1 | 458718 | *USP9Y, UMN2001* |
| NW_017189685.1 | 386696 | *DBY, DDX3Y, UTY* |
| NW_017189885.1 | 194156 | *ZFY* |

**Table 3.** Codes and SRA accession numbers of WGS data used for Fig. 1.

| **code [1]** | **our code** | **SRA accession** | **breed** | **haplogroup** | **reference** |
| --- | --- | --- | --- | --- | --- |
| BDCH03 | BDBLB03 | SRX2982563 | Black Bengal | Y2B | [1] |
| CNNCH04 | CNSHD04 | SRX2982510 | Shandong Yimeng black | Y1AB | [1] |
| CNNCH06 | CNXIN06 | SRX2982522 | Xinjiang | Y1AB | [1] |
| CNNCH14 | CNQIN14 | SRX2982537 | Qinghai | Y1AB | [1] |
| CNNCH15 | CNQIN15 | SRX2982502 | Qinghai | Y1AB | [1] |
| CNNCH16 | CNQIN16 | SRX2982503 | Qinghai | Y1AB | [1] |
| CNNCH17 | CNQIN17 | SRX2982504 | Qinghai | Y1AB | [1] |
| CNNCH18 | CNQIN18 | SRX2982511 | Qinghai | Y1AB | [1] |
| CNNCH19 | CNQIN19 | SRX2982507 | Qinghai | Y1AB | [1] |
| CNNCH20 | CNQIN20 | SRX2982487 | Qinghai | Y1AB | [1] |
| CNSCH09 | CNRAO09 | SRX2982486 | Raoshan Henan | Y1AA | [1] |
| CNTCH01 | CNTIB01 | SRX2982520 | Tibetan | Y1AB | [1] |
| CNTCH02 | CNTIB02 | SRX2982521 | Tibetan | Y2B | [1] |
| CNTCH03 | CNTIB03 | SRX2982513 | Tibetan | Y2B | [1] |
| CNTCH04 | CNTIB04 | SRX2982514 | Tibetan | Y1AB | [1] |
| CNTCH05 | CNTIB05 | SRX2982515 | Tibetan | Y1AB | [1] |
| FRCH06 | FRPTV06 | SRX2982542 | Poitou | Y1AB | [1] |
| NLCH01 | NLLNR01 | SRX2982482 | Dutch Landrace | Y1B | [1] |
| NLCH03 | NLDSA03 | SRX2982490: | Dutch Saanen | Y1B | [1] |
| PKCH05 | PKBBL05 | SRX2982568 | Beetal Black | Y2A | [1] |
| SDCH03 | AFSD03 | SRX5417024 | Sudan | Y2A | [1] |
| SDCH05 | AFSD05 | SRX5417026 | Sudan | Y2A | [1] |
| FRCH05 | FRALP05 | SRX2982481 | Alpine | Y1B | [1] |
|  | KOSAA04 | ERX2360419 | Korean Saanen | Y1B2 | [2] |
|  | KOSAA05 | ERX2360420 | Korean Saanen | Y1B2 | [2] |
|  | KOSAA08 | ERX2360423 | Korean Saanen | Y1B | [2] |
|  | KOSAA09 | ERX2360424 | Korean Saanen | Y1B | [2] |
|  | KOSAA10 | ERX2360425 | Korean Saanen | Y1B2 | [2] |
|  | KOKOR01 | ERX2360430 | Korean Saanen | Y1B | [2] |
|  | KOKOR02 | ERX2360431 | Korean Saanen | Y1AA | [2] |
|  | KOKOR08 | ERX2360437 | Korean Saanen | Y1B | [2] |
|  | KOKOR10 | ERX2360439 | Korean Saanen | Y1B | [2] |
|  | KOKOR11 | ERX2360440 | Korean Saanen | Y1B | [2] |
|  | KOKOR281 | ERX2360442 | Korean Saanen | Y1B | [2] |
|  | KOKOR282 | ERX2360443 | Korean Saanen | Y1B | [2] |
|  | KOKOR04 | ERX2360445 | Korean Saanen | Y2B | [2] |
| IRCH04 | IRA04 | ERX286352 | Iran | Y2A | [3] |
| IRCH06 | IRA06 | ERX286345 | Iran | Y2A | [3] |
| IRCH10 | IRA10 | ERX286355 | Iran | Y1AA | [3] |
| IRCH14 | IRA14 | ERX286363 | Iran | Y2A | [3] |
| IRCH16 | IRA16 | ERX286347 | Iran | Y2A | [3] |
| IRCH19 | IRA19 | ERX313213 | Iran | Y1AB | [3] |
| IRCH20 | IRA20 | ERX313216 | Iran | Y1AB | [3] |
| MCCH01 | MOR01 | ERX194212 | Moroccan | Y2A | [3] |
| MCCH02 | MOR02 | ERX194211 | Moroccan | Y2A | [3] |
| MCCH03 | MOR03 | ERX204035 | Moroccan | Y2A | [3] |
| MCCH04 | MOR04 | ERX204033 | Moroccan | Y2A | [3] |
| MCCH05 | MOR05 | ERX204019 | Moroccan | Y2A | [3] |
| MCCH08 | MOR08 | ERX204026 | Moroccan | Y2A | [3] |
| MCCH10 | MOR10 | ERX204034 | Moroccan | Y2A | [3] |
| MCCH11 | MOR11 | ERX204027 | Moroccan | Y2A | [3] |
| MCCH12 | MOR12 | ERX207043 | Moroccan | Y2A | [3] |
| MCCH15 | MOR15 | ERX208812 | Moroccan | Y2A | [3] |
| MCCH17 | MOR17 | ERX220684 | Moroccan | Y2A | [3] |
| MCCH23 | MOR23 | ERX272615 | Moroccan | Y1AB | [3] |
| MCCH24 | MOR24 | ERX286410 | Moroccan | Y1AB | [3] |
| MCCH25 | MOR25 | ERX286408 | Moroccan | Y2A | [3] |
| MCCH26 | MOR26 | ERX286419 | Moroccan | Y2A | [3] |
| MCCH28 | MOR28 | ERX286420 | Moroccan | Y2A | [3] |
| MCCH29 | MOR29 | ERX286403 | Moroccan | Y1AB | [3] |
| MCCH31 | MOR31 | ERX288658 | Moroccan | Y2A | [3] |
| MCCH33 | MOR33 | ERX288657 | Moroccan | Y2A | [3] |
| MCCH37 | MOR37 | ERX288652 | Moroccan | Y2A | [3] |
| MCCH38 | MOR38 | ERX288656 | Moroccan | Y1AB | [3] |
| MCCH39 | MOR39 | ERX288926 | Moroccan | Y2A | [3] |
| MCCH40 | MOR40 | ERX291909 | Moroccan | Y2A | [3] |
| MCCH41 | MOR41 | ERX306478 | Moroccan | Y2A | [3] |
| MCCH42 | MOR42 | ERX306471 | Moroccan | Y2A | [3] |
| MCCH43 | MOR43 | ERX313276 | Moroccan | Y2A | [3] |
| MCCH44 | MOR44 | ERX286409 | Moroccan | Y2A | [3] |

[1] Z. Zheng & Y. Jiang, unpublished
[2] Kim et al. (2019) doi: 10.3389/fgene.2019.00699
[3] Alberto et al. (2018) doi: 10.1038/s41467-018-03206-y

### 2.2 Haplogroups in 132 goat breeds

We differentiating the major Y-chromosomal haplogroups Y1AA, Y1AB, Y1B, Y2A and Y2B for 1720 domestic goats from 132 breeds by combining data from several sources as detailed per breed in Table 4:

1. From the goat panel collected for the Econogene project, DNA samples of 353 male goats from 38 European or southwestern Asian breeds were analyzed by PCR amplification and dideoxy-sequencing of segments within the *DBY, SRY* and *ZFY* genes (Lenstra and Econogene Consortium, 2005) as described previously for bovine samples (Nijman et al., 2008; Edwards et al., 2011) and using the primers listed in Table 5.
2. We used published data for 90 goats from 5 Portuguese breeds or from Morocco (Pereira et al., 2009), for 211 goats from 10 Asian breeds (Waki et al., 2015), for 181 goats from 8 Turkish breeds (Cinar Kul et al., 2015) and for 270 goats from 26 Spanish or Swiss breeds (Vidal et al., 2017).
3. 362 DNA samples (57 breeds) from several sources, including the AdaptMap panel (Colli et al., 2018) were genotyped by the KASP assay (Table 4).
4. For 90 goats from 15 breeds genotypes were obtained by Blast searching of the ShortRead Archive (SRA).

**Table 4.**
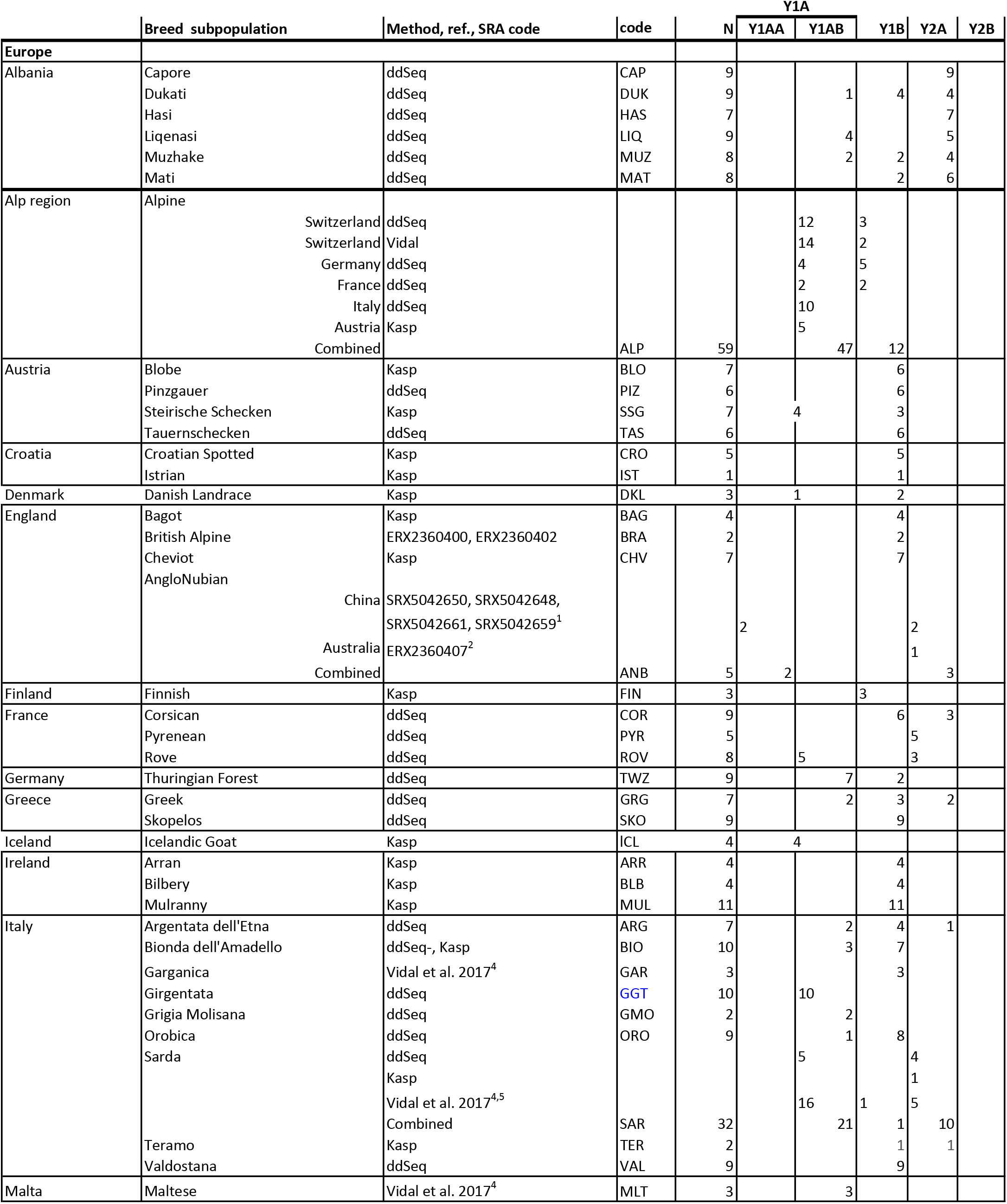
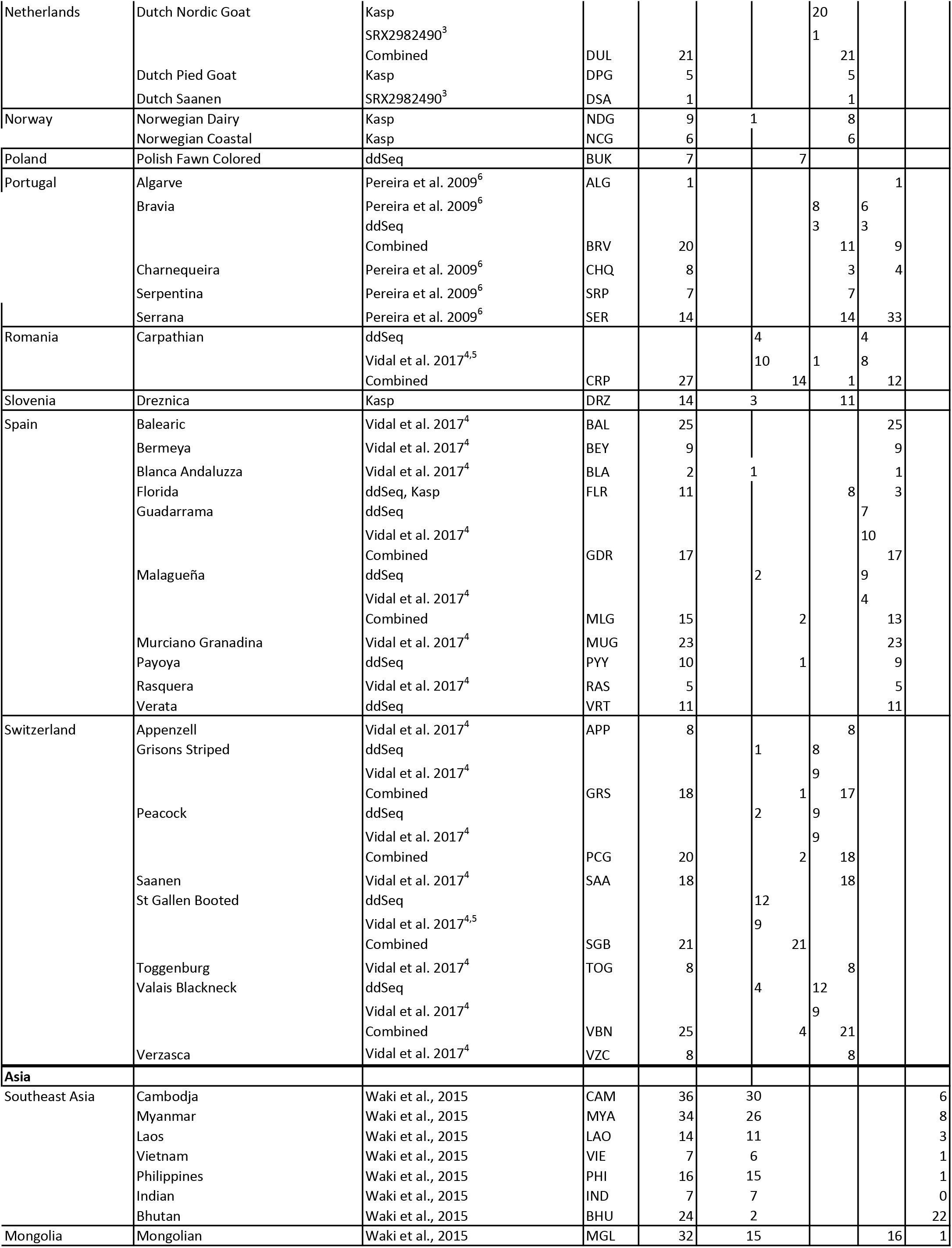
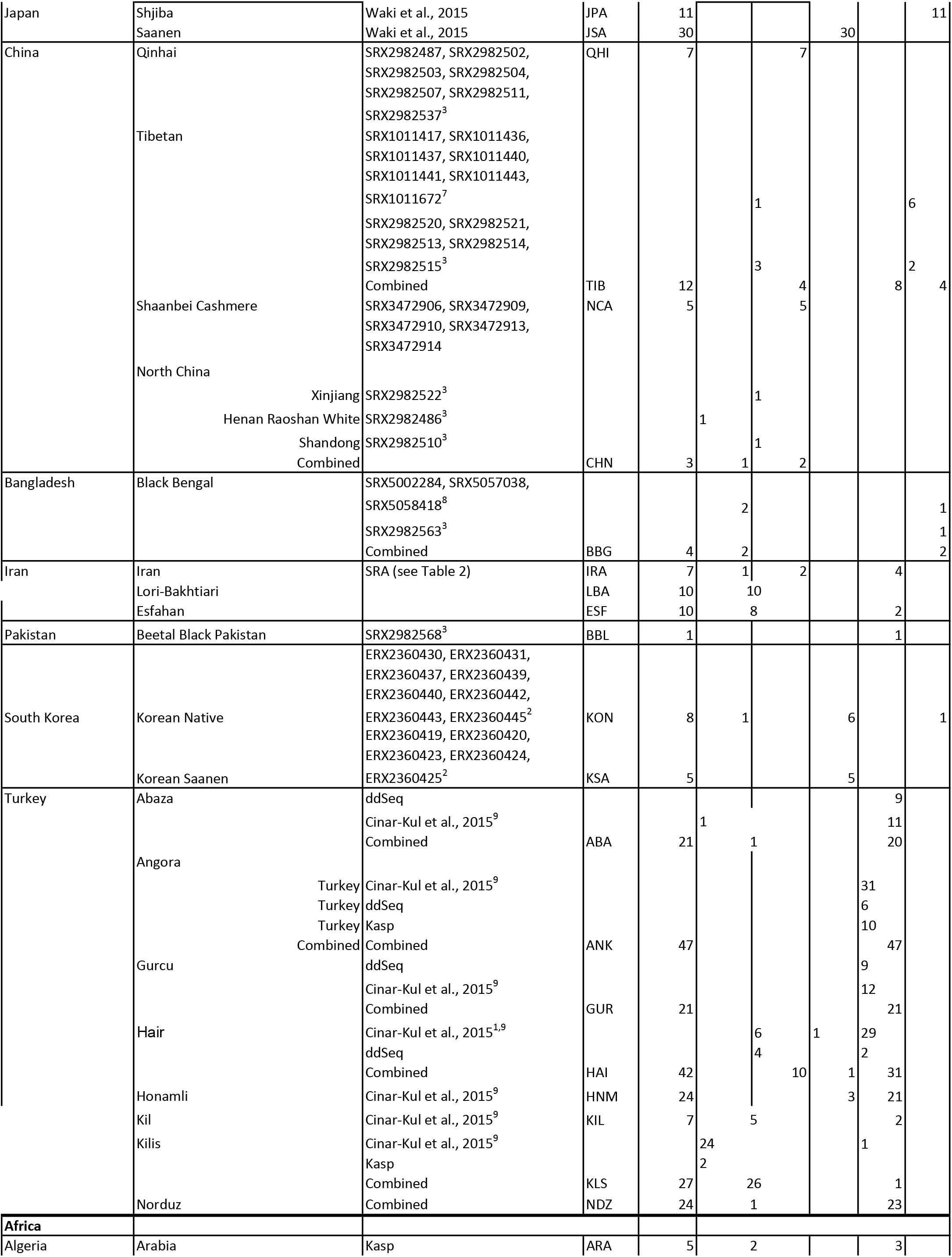
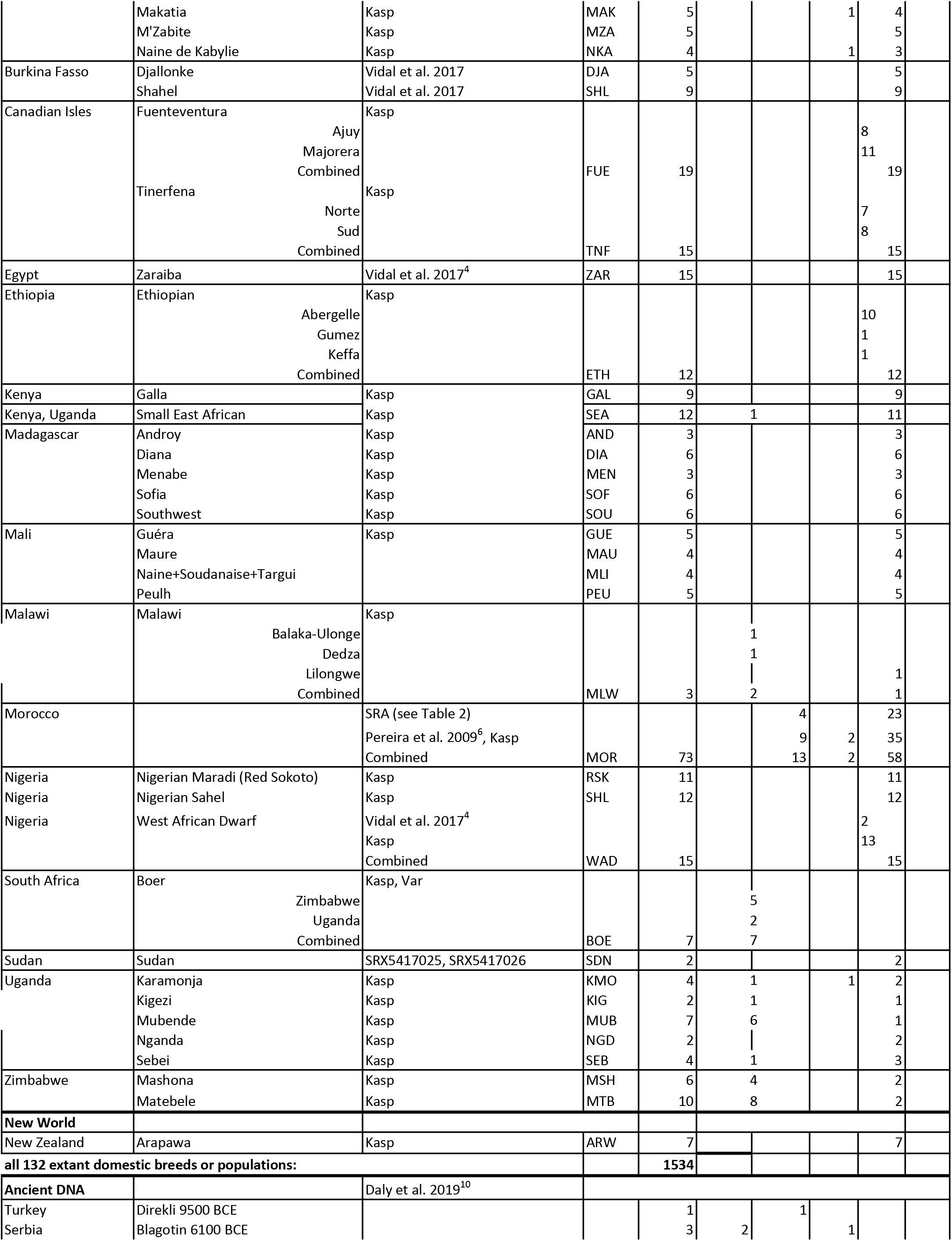
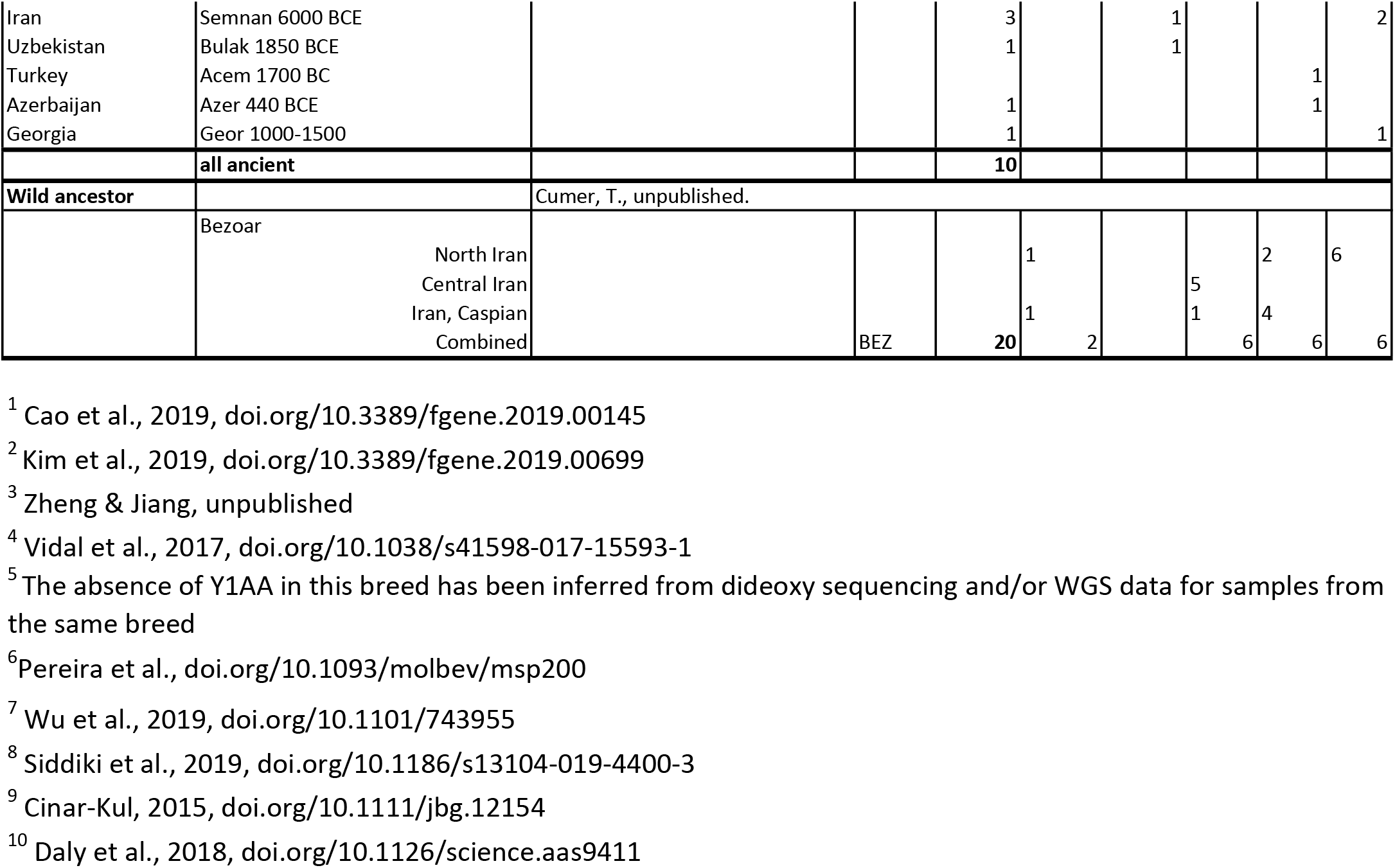
Haplogroup distributions in breeds or populations, in ancient DNA and in the bezoar. Kasp, Kasp (Kompetitive allele specific PCR assay); ddSeq, dideoxy sequencing. Codes starting with ERX or SRX codes are from the Short Reads Archive

**Table 5.**
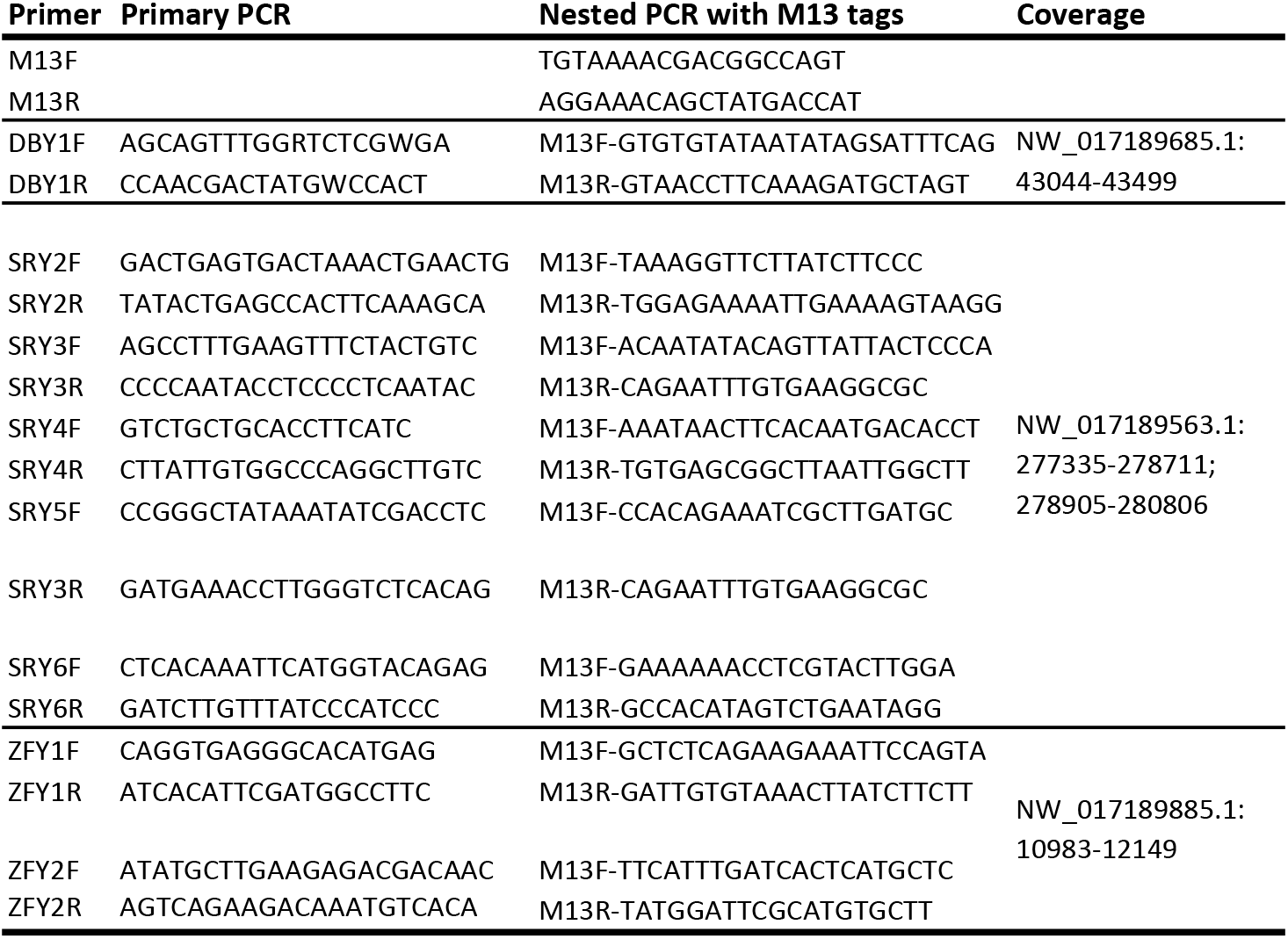
Primers for amplification and dideoxy sequencing of DBY, SRY and ZFY fragments and genomic coordinates of sequence covered by the sequence reads excluding the primer sequences.

For samples collected by Cinar-Kul (2015) and Vidal et al. (2017) or analyzed by KASP before genomic data became available, Y1AA and Y1AB have been both scored as Y1A haplotype. Several of these samples did not contain Y1A or belonged to breeds for which additional data were available (Table 3). However, for 31 breeds we only have the Y1A frequency. Likewise, Vidal et al. (2017) and the Kasp assay did not differentiate Y2A and Y2B. Since sequence data indicate that Y2B does not occur in modern domestic goats from Europe, Anatolia and Iran, we assigned all Y2 scores in European and African goats to Y2A.

Genotypes were extracted from WGS data for ancient DNA samples (Daly et al. 2018) and for bezoar samples (Alberto et al., 2018).

## 3. Results and Discussion

The four selected scaffolds (Table 1) cover together 1,567,760 bp of the male-specific part of the caprine Y-chromosome and contain the SNPs that in previous work defined the major haplotypes Y1A, Y1B, Y2A and Y2B (Table 1). A phylogenetic tree of the Y-chromosomes of 70 goats (Fig. 1) shows five major haplogroups. Three haplogroups contains the previously identified haplotypes Y1B, Y2A and Y2B, whereas the Y1A haplotypes belong to either haplogroups Y1AA and Y1AB. In this panel we did not find goats with the local Y1C and Y2C sequences (Table 1).

**Fig. 1.**
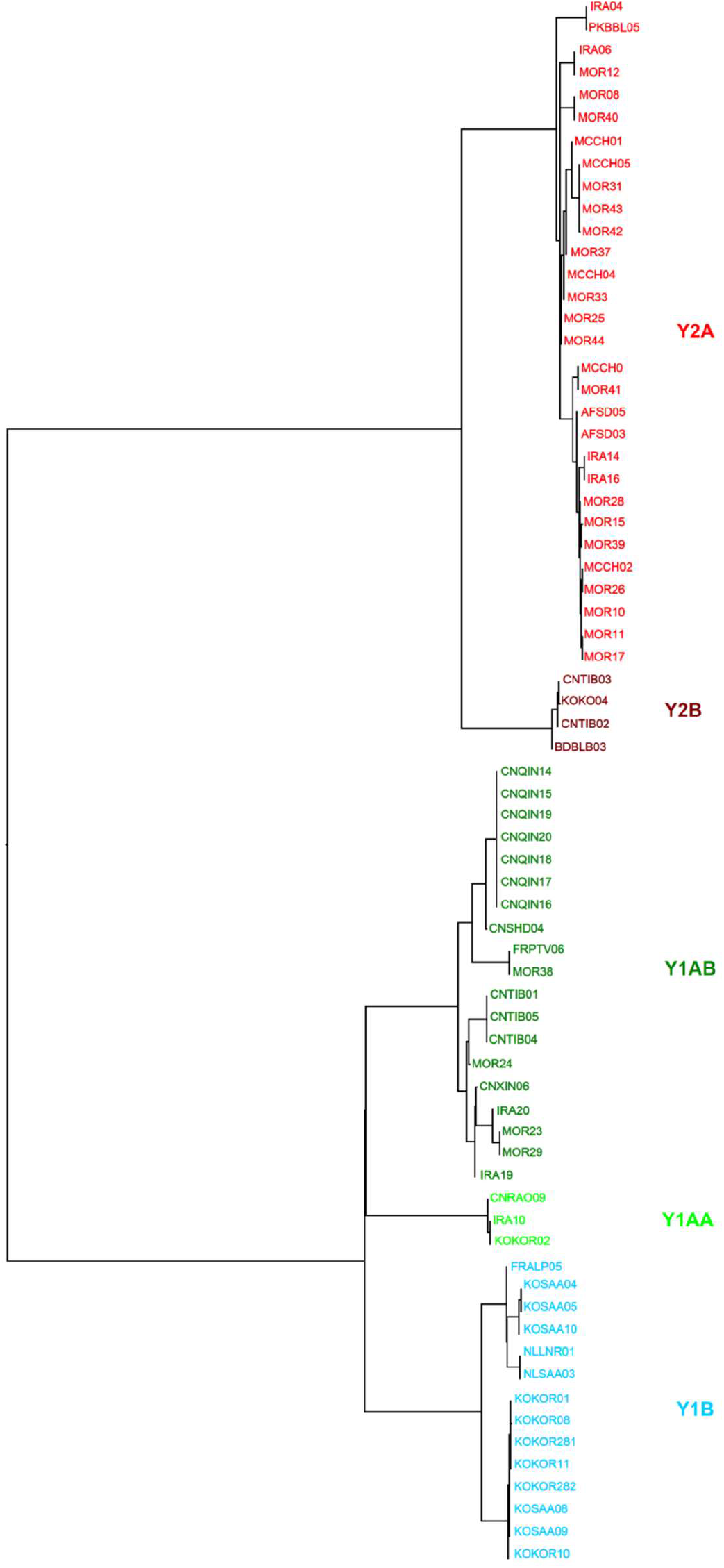
Neighbor-joining tree of allele-sharing distances calculated on the basis of 2350 hemizygous male-specific SNPs extracted from WGS of 70 male goats (Table 3) and the markhor sequence as an outgroup. AFSD, Sudan; BDBLB, Black Bengal; CNQIN, Qinghai; CNRAO, Henan Raoshan White; CNSHD, Shandong Yimeng Black; CNTIB, Tibetan; CNXIN, Xinjiang; FRALP, French Alpine; FRPTV, Poitou; IRA, Iran; KOKOR, Korea native; KOSAA, Korean Saanen; MOR, Morocco; NLLNR, Dutch Landrace; NLDSA, Dutch Saanen; PKBBL, Pakistan Beetal Black.

**Fig. 2.**
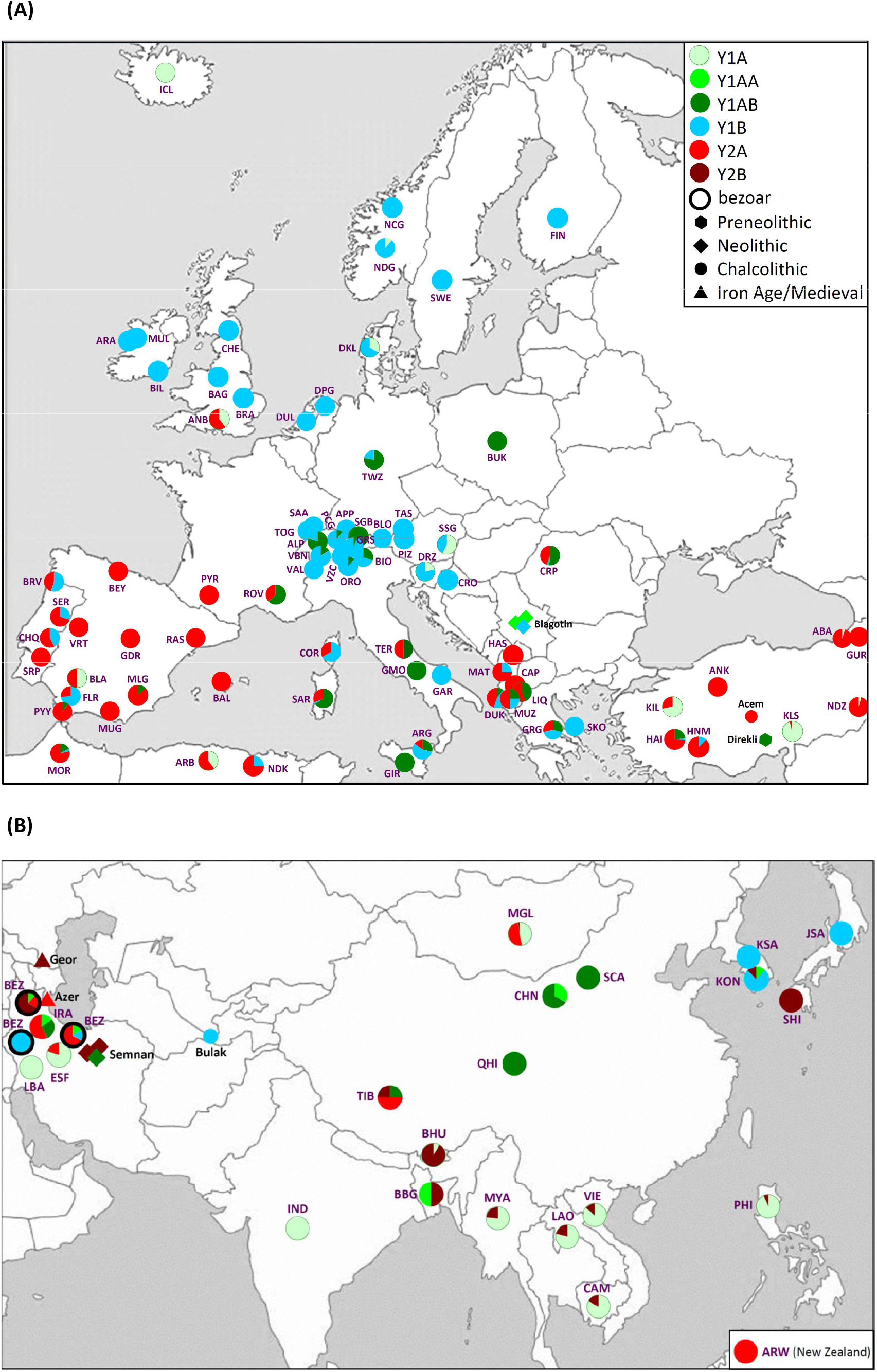
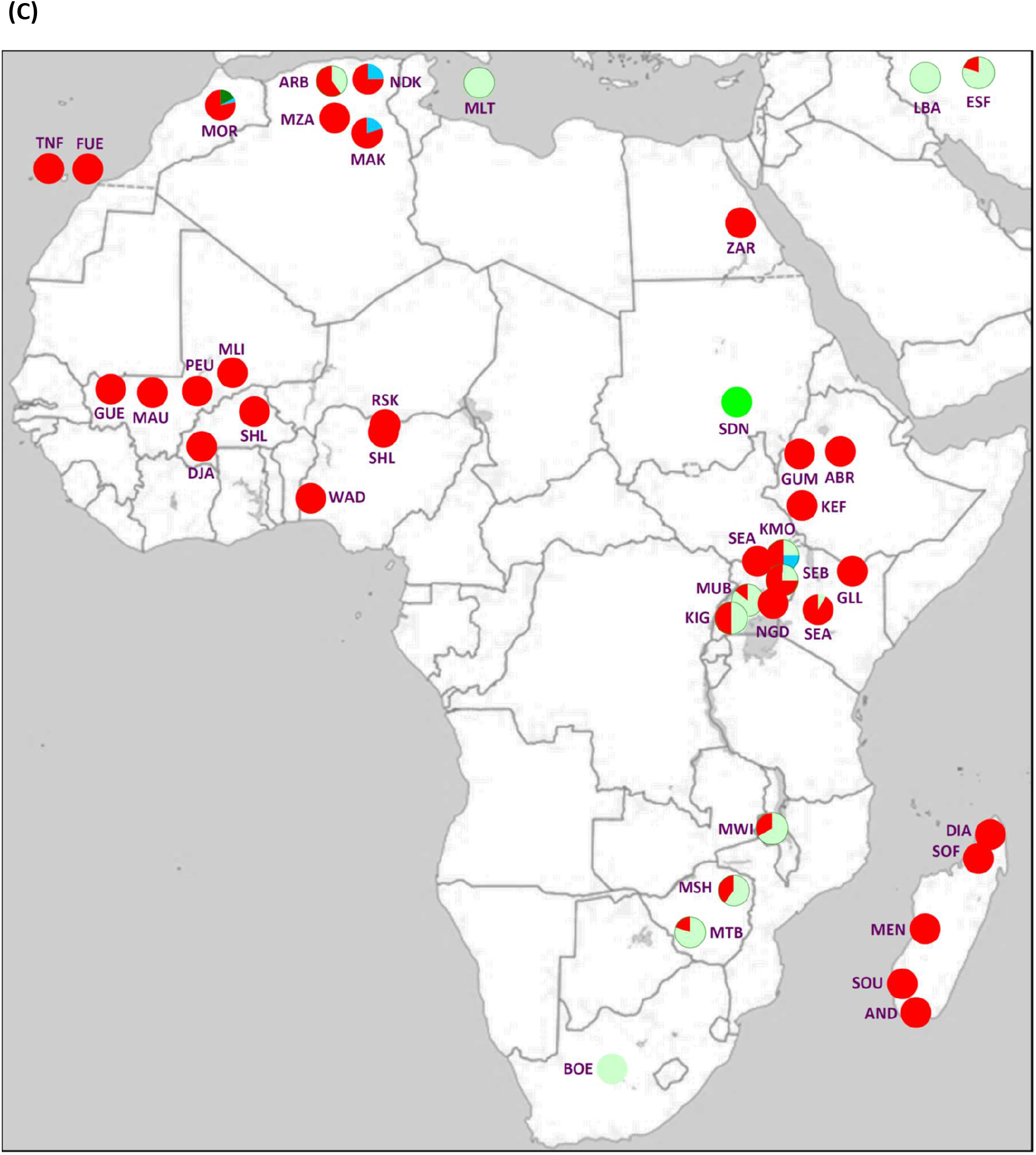
Haplogroup distributions of **(A)** European breeds; **(B)** Asian breeds; **(C)** African breeds; **(A, B)**, European and Asian ancient DNA samples; and **(B)** Iranian bezoars. Breeds represented by a single goat are not plotted or are combined with other breeds from the same country as indicated in Table 4. Breed codes: ABA, Abaza; ALP, Alpine; ANB, AngloNubian; AND, Androy; ANK, Angora;Ankara; APP, Appenzell; ARA, Arabia; ARG, Argentata dell’Etna; ARR, Arran; ARW, Arapawa; BAG, Bagot; BAL, Balearic; BBG, Black Bengal; BEY, Bermeya; BHU, Bhutan; BIO, Bionda dell’Amadello; BLA, Blanca Andaluzza; BLB, Bilbery; BLO, Blobe; BOE, Boer; BRA, British Alpine; BRV, Bravia; BUK, Polish Fawn Colored; CAM, Cambodja; CAP, Capore; CHN, North China (Xinjiang, Henan Raoshan White, Shandong); CHQ, Charnequeira; CHV, Cheviot; COR, Corsican; CRO, Croatian Spotted; CRP, Carpathian; DIA, Diana; DJA, Djallonke; DKL, Danish Landrace; DPG, Dutch Pied Goat; DRZ, Dreznica; DUK, Dukati; DUL, Dutch Nordic Goat; ESF, Esfahan; ETH, Ethiopian (Abergelle, Gumez, Keffa); FIN, Finnish; FLR, Florida; FUE, Fuenteventura (Ajuy, Majorera); GAL, Galla; GAR, Garganica; GDR, Guadarrama; GGT, Girgentata; GMO, Grigia Molisana; GRG, Greek; GRS, Grisons Striped; GUE, Guéra; GUR, Gurcu; HAI, Hair; HAS, Hasi; HNM, Honamli; IND, Indian; IRA, Iranian; JPA, Shjiba; JSA, Japanese Saanen; KIG, Kigezi; KIL, Kil; KLS, Kilis; KMO, Karamonja; KON, Korean Native; KSA, Korean Saanen; LAO, Laos; LBA, Lori-Bakhtiari; lCL, Icelandic; LIQ, Liqenasi; MAK, Makatia; MAT, Mati; MAU, Maure; MEN, Menabe; MGL, Mongolian; MLG, Malagueña; MLI, Naine, Soudanaise Targui; MLT, Maltese; MLW, Malawi (Balaka-Ulonge, Dedza;Lilongwe; MOR, Moroccan; MSH, Mashona; MTB, Matebele; MUB, Mubende; MUG, Murciano Granadina; MUL, Mulranny; MUZ, Muzhake; MYA, Myanmar; MZA, M’Zabite; NCG, Norwegian Coastal; NDG, Norwegian Dairy; NDZ, Norduz; NGD, Nganda; NKA, Naine de Kabylie; ORO, Orobica; PCG, Peacock; PEU, Peulh; PHI, Philippines; PIZ, Pinzgauer; PYR, Pyrenean; PYY, Payoya; QHI, Qinhai; RAS, Rasquera; ROV, Rove; RSK, Nigerian Maradi (Red Sokoto); SAA, Saanen; SAR, Sarda; SCA, Shaanbe Cashmere; SDN, Sudan; SEA, Small East African (Kenya, Uganda); SEB, Sebei; SRP, Serpentina; SER, Serrana; SGB, St Gallen Booted; SHL, Shahel; SHL, Nigerian Sahel; SKO, Skopelos; SOF, Sofia; SOU, Southwest; SSG, Steirische Schecken; TAS, Tauernschecken; TER, Teramo; TIB, Tibetan; TNF, Tinerfena (Norte, Sud); TOG, Toggenburg; TWZ, Thuringian Forest; VAL, Valdostana; VBN, Valais Blackneck; VIE, Vietnam; VRT, Verata; VZC, Verzasca; WAD, West African Dwarf; ZAR, Zaraiba.

Remarkably, with the exception of Y1B the haplogroups have all been found in Iranian bezoar samples (Fig. 3, Table 4), whereas all haplogroups, including Y1B, were detected in ancient goat samples.

Geographic plots of haplogroup frequencies show a considerable spatial differentiation, which resonates the strong phylogeography displayed by autosomal SNPs (Colli et al., 2018), but is in clear contrast with the phylogenetic structure displayed by mtDNA haplogroups (Luikart et al., 2001; Naderi et al., 2007, 2008; Zhao et al., 2014b, 2014a). Most likely, by a series of bottlenecks in the male lineage subcontinents have different major haplogroups, while none has a global-wide coverage:

- Haplogroup Y2B is absent in Europe, Africa and west Asia, but became a major haplotype in east Asia and southeast Asian, where Y2A is not represented. In contrast, it is observed in ancient goat from Medieval Georgia and Neolithic Iran (ca. 6,000 BCE), supporting an origin of east Asian goat from regions east of Zagros Mountains.
- Y2A is in northern and central Europe only found in the crossbred AngloNubian (see below), but is the predominant haplogroup in central, eastern and southern Africa.
- Haplogroup Y1B is predominant in northern Europe, but outside Europe and North African has only been found in one Karamonja animal, in the Korean native breed and in exported Saanen populations. The different Y2A and Y1B frequencies in north-central vs southern Europe may reflect the Neolithic migrations following the Danube and the Mediterranean routes, respectively (Cymbron et al., 2005; Tresset and Vigne, 2007; Rivollat et al., 2015) with the strongest bottlenecks along the northernmost migrations.
- Y1AA is sporadic in Europea and has been observed also in Neolithic Serbia (ca. 6,000 BCE), but is present in Asia (Fig. 3).

Deviations from the general pattern may very well reflect major introgressions. A well-known example is the Anglo-Nubian, which originated in England by crossing Indian or African imported goats with local breeds and is in our panel the only northern-European breed carrying Y2A.

There were two out-of-range findings of Y1B, in the Uganda Karamonja and in the Korean native breeds. Because of the popularity of Swiss dairy goats in both Uganda (NAADS, 2005) and Korea (Kim et al., 2019), crossbreeding again is the most likely explanation.

We conclude that the Y-chromosomal haplotype distribution reveals expansion and crossbreeding events, as observed also for the human paternal lineages (Poznik et al., 2016; Batini and Jobling, 2017), and illustrates the power of Y-chromosomal markers for inferring the genetic origin of mammalian populations.

## Authors’ contributions

IJN, PAM and JAL designed the study; IJN carried out and analyzed the ABI sequencing; TF, BD, BR, ZZ and YJ analyzed the WGS data; TC and FPo supplied the bezoar genotypes; KGD and DGB supplied the aDNA data; BR supplied most of the African samples for KASP genotyping; BB, VAB, TB, SC, VC-C, AdS, JK, NK, AM, RM, FPe MS, AS, JS and JT collected material or data for other breeds; JAL performed the downstream analysis and wrote the first draft; and KGD, AM, FPe, MS and PAM contributed to the text.

## Acknowldegements

This study has benefited from an interaction with Vargoats consortium (http://www.goatgenome.org/vargoats.html).

## Availability of data

The SRA data are in the public domain.

## Members of the Econogene Consortium

**Albania:** Hoda Anila, Dobi Petrit, Dep. Animal Production, Fac. Agriculture, Tirana; **Austria:** Baumung Roswitha, Univ. Natural Resources Applied Life Sciences, Vienna; **Belgium:** Baret Philippe, Fadlaoui Aziz, Dep. Animal Genetics, Univ. Cath. Louvain, Louvain-la-Neuve; **Cyprus:** Papachristoforou Christos, Agricult. Research Instit., Nicosia; **Egypt:** El-Barody M.A.A., Animal Production Dep., Fac. Agriculture, Minia Univ. **France:** Taberlet Pierre, England Phillip, Luikart Gordon, Beja-Pereira Albano, Zundel Stéphanie, Laboratoire d’Écologie Alpine (LECA), Univ. Joseph Fourier, Grenoble; Trommetter Michel, Inst. Recherche Agronomique, Unité d’Economie Sociologie Rurales, Grenoble; **Germany:** Erhardt Georg, Brandt Horst, Ibeagha-Awemu Eveline, Lühken, Gesine, Daniela Krugmann, Prinzenberg Eva-Maria, Lipsky Shirin, Gutscher Katja, Peter Christina, Inst. Tierzucht Haustiergenetik, Justus-Liebig Univ. Giessen; Roosen Jutta, Bertaglia Marco, Dep. Food Economics Consumption Studies, Univ. Kiel; **Greece:** Georgoudis Andreas, Al Tarrayrah, Jamil, Kliambas Georgios, Kutita Olga, Karetsou Katerina, Aristotle Univ. Thessaloniki, Fac. Agriculture, Thessaloniki; Ligda Christina, National Agric. Research Foundation, Thessaloniki; **Hungary:** Istvan Anton, Fesus Lazlo, Research Inst. Animal Breeding Nutrition, Dep. Genetics, Herceghalom; **Italy:** Ajmone-Marsan Paolo, Canali Gabriele, Milanesi Elisabetta, Pellecchia Marco, Università Cattolica S. Cuore, Piacenza; Carta Antonello, Sechi Tiziana, Ist. Zootecnico Caseario Sardegna, Sassari; Cicogna Mario, Fornarelli Francesca, Giovenzana Stefano, Marilli Marta, Ist. Zootecnia Generale, Univ. Studi di Milano; Marletta Donata, Bordonaro S., D’Urso Giuseppe, Dip. Scienze Agronomiche Produzioni Animali, Univ. Studi Catania; Pilla Fabio, D’Andrea Mariasilvia, Dip. Scienze Animali Vegetali Ambiente, Univ. Molise, Campobasso; Valentini Alessio, Cappuccio Irene, Pariset Lorraine, Dip. Produzioni Animali, Univ. Tuscia, Viterbo; **Jordan:** Abo-Shehada Mahamoud, Fac. Veterinary Medicine, Jordan Univ. Science Technology; **Netherlands:** Lenstra Johannes A., Nijman, Isaäc J., Van Cann, Lisette M., Fac. Veterinary Medicine, Utrecht Univ.; **Poland:** Niznikowski Roman, Dominik Popielarczyk, Strzelec Ewa, Dep. Sheep Goat Breeding, Warsaw Agricultural Univ.; **Romania:** Vlaic Augustin, Dep. Animal Genetics, Faculty Zootechnics, Univ. Cluj-Napoca; **Spain:** Dunner Susana, Canon Javier, Cortes Oscar, Garcia David, Univ. Complutense Madrid; **Switzerland:** Caloz Régis, EPFL, Lausanne; Obexer-Ruff Gabriela, Marie-Louise Glowatzki, Institut für Genetik Ernährung und Haltung von Haustieren, Universität Bern; **Turkey:** Ertugrul Okan, Veterinary Fac., Ankara Univ.; Togan Inci, Koban Evren, Dep. Biology, Middle East Technical Univ., Ankara; **UK:** Bruford Mike, Perez Trinidad, Juma Gabriela, School Biosciences, Cardiff Univ.; Hewitt Godfrey, Dalamitra Stella, Wiskin Louise, Taylor Martin, Biological Sciences, Univ. East Anglia, Norwich; Jones Sam, The Sheep Trust, Scarpa Riccardo, Environment Dep., Univ. York

